# Characterization of a regulatory T cells molecular meta-signature identifies the pro-enkephalin gene as a novel marker in mice

**DOI:** 10.1101/638072

**Authors:** Nicolas Aubert, Benoit L. Salomon, Gilles Marodon

**Author notes:** correspondence to, Sorbonne University, CIMI-PARIS, 91 Bd de l’Hôpital, 75013 PARIS.

## Abstract

Regulatory T cells (Treg) are crucial in the proper balance of the immune system. A better characterization of Treg-specific genes should extend our knowledge on their complex biology. However, to date there is no consensual Treg signature in the literature. Here, we extracted a molecular Treg meta-signature relative to CD4^+^ conventional T cell from 8 different but comparable publicly available microarray datasets. We confirmed the validity of our result using the much larger but less stringent Immuno-Navigator database. However, many genes of the Treg meta-signature were also expressed at the protein level by other immune cell subsets, as assessed by mass cytometry, with the noticeable exceptions of *Il2ra*, *Ctla4*, and *Tnfrsf9. S*urprisingly, the proenkephalin (*Penk*) gene was a prominent member of this restricted Treg meta-signature. Further analysis of public datasets and of our own RNA sequencing experiments confirms that *Penk* was over expressed by Treg in various murine tissues, including thymic Treg. Interestingly, *Penk* expression was increased in intra tumoral Treg whereas it was down modulated in the central nervous system of mice suffering from EAE. Finally, we propose a mechanistic model linking TNFR signaling and the transcription factor *Batf* in the regulation of *Penk* expression in Treg. Altogether, our results provide the first Treg meta-signature in mice and identifies *Penk* as a novel and unexpected Treg marker.

## Introduction

Proper number and function of regulatory T cells (Treg) are essential for a well-balanced immune system: too few of these cells leads to autoimmunity and too much prevents an efficient immune response, with harmful consequences for anti-tumor immunity, for instance. Treg are a subset of CD4+ T cells that express the transcription factor (TF) Foxp3 and the alpha chain of the interleukin-2 receptor CD25, both indispensable for suppressive functions and proper homeostasis. Probably the best example illustrating the crucial role for Foxp3-expressing cells in the homeostasis of the immune system is given by the lethal auto immune syndrome seen in patients bearing mutations in the *FOXP3* gene, the IPEX syndrome (1). Like most CD4^+^ T cells, Treg are generated in the thymus upon MHC-driven selection based on affinity of the T cell receptor for self antigens (2). A sizable proportion of those cells are also induced in the periphery (pTreg) from CD4+ T cells precursors, but those pTreg are reported not to express Helios or Neuropilin-1, contrary to thymic-derived Treg (tTreg) that are positive for those markers (3). Known pTreg inducers in the periphery can be byproducts of bacterial metabolism (4,5) but it is likely that inflammation *per se* is a main driver for pTreg differentiation (6). Thus, finding ways of manipulating Treg for therapeutic purposes in the auto immunity field has become a major endeavor for immunologists worldwide. Moreover, recent results linking the presence of Treg to a bad prognostic in some cancers extended their potential clinical applications from autoimmunity to cancer immunotherapy (7). In the case of cancer, one would want to get rid of Treg to wake up a dim immune response to tumors. An example of this powerful approach has been recently illustrated by Treg-depleting CTLA-4 or CD25-specific mAb (8,9). However, this weak response to tumors is part of a natural tolerance process, preventing the immune system to attack self-tissues. Thus, breaking immune tolerance by removing Treg is not without consequences on the integrity of healthy tissues. This is illustrated by studies showing that Ipilimumab (anti-CTLA-4), a powerful anti-cancer drug affecting Treg, indeed help the immune system to fight tumors at the expense of a generalized auto immunity in treated patients (10). A better knowledge of Treg biology will be crucial for preserving therapeutic efficacy without severe adverse events.

This knowledge has been mostly collected from mice due to their ease of use, their versatility and the thousands of genetic models available to answer mechanistic questions. To that end, hundreds of investigators have pursued the quest for specific Treg markers, revealing molecules and pathways that can be targeted by monoclonal antibodies or pharmacological compounds. However, there are still confusions about these specific markers, since the comparisons are often made across unrelated studies employing different technologies. Moreover, a characteristic of Treg is their ability to adapt to the cellular environment in which they are present. This mechanism, referred as to effector class control few years ago (11), deeply affect cell surface phenotype and thus presumably, gene expression patterns. In addition, there are evidences that pTreg, contrary to tTreg, might be relatively unstable (12), meaning that pTreg may acquire some pro-inflammatory effector functions. Furthermore, it appears that tissue Treg also differ from Treg of lymphoid organs at the molecular level, with the TF BACH2 repressing expression of tissue-specific genes (13) whereas the TF BATF induced expression of these genes (14). One can immediately realize that providing a common definition of Treg across this diversity of phenotype and function represents a difficult challenge. Furthermore, Treg specific markers should ideally mark Treg only or should be minimally expressed by other cells of the immune system or other non-immune cells. This has been an overlooked issue since most of the so-called “Treg signatures” are established relative to CD4^+^ T conventional cells (Tconv), and not looking outside of Tconv (not mentioning major phenotypic differences used to define Treg and Tconv). Safety and efficacy of Treg-based therapies will surely benefit from targeting molecules and/or pathways truly specific to Treg.

In an attempt to resolve some of these issues, we reasoned that digging out a Treg meta-signature (TMS) from available datasets comparing well-defined Treg and Tconv should lead to a more robust signature than studies taken separately. As a starting point, we decided to focus our investigation in Mus Musculus, because many databases and tools are available in mice. We also focused our analysis on resting Treg freshly isolated *ex vivo*, taken from non inflammatory lymphoid organs, to avoid bias due to T cell activation and/or tissue localization. This naive approach led to the first description of a “universal” Treg signature, which includes several known Treg specific markers but also new ones, such as the pro-enkephalin gene *Penk*, indicating a function for the endogenous opioid pathway in immunoregulation.

## Methods

### Extraction of Treg meta-signatures

The datasets used were selected based on a “Treg* AND (Tconv* OR Teff*) AND Mus Musculus” search in the GEO dataset web site (https://www.ncbi.nlm.nih.gov/gds). GEO datasets were manually inspected for inclusion of studies comparing fresh Treg with fresh Tconv from lymphoid organs. Only GSE17580 (15), GSE24210 (16), GSE37532 (17,18), GSE40685 (19), GSE42021 (20), GSE7852 (21), GSE50096 (22), and GSE15907 (ImmGen Project (23)) were selected and a list of genes significantly up regulated in Treg compared to Tconv was determined for each dataset using GEO2R with an adjusted p.value cutoff of 0.05 (False Discovery Rate). The commonality within gene sets was determined using the online tool from the Bioinformatics Evolutionary Genomics department form the Ghent University (http://bioinformatics.psb.ugent.be/webtools/Venn/). The Treg signature from Immuno Navigator (https://genomics.virus.kyoto-u.ac.jp/immuno-navigator/?) was generated by extracting 634 CD4 T cell samples and 240 Treg samples from the database. Differential gene expression was determined using Qlucore v3.4 with optimal variance set at 0.5 and False Discovery Rate <0.05. All but one dataset (GSE37532) where the Affymetrix Mouse Gene 1.0 ST Array was used, were generated using the Affymetrix Mouse Genome 430 2.0 Array.

### Analysis of the TMS

All network analysis and visual representations were performed with Cytoscape v3.7 (24) (https://cytoscape.org). For enrichment analysis, we used ClueGO v2.5.5 and CluePedia 1.5.5 that integrates several ontology and pathway databases and creates a functionally organized network of GO/pathway terms (25,26). To represent PPI networks, we used the GeneMania (27) or STRING (28) Cytoscape applications, with embedded enrichment analysis. Expression of *Penk* in various tissues is represented by the graphical tool embedded in GEO2R, the Jamovi software (29) or Prism v8.3 (Graphpad). Putative regulators of *Penk* mRNA in mice was generated with Ingenuity Pathway Analysis (Qiagen)

### Mice

All mice were on a C57Bl/6J background. Foxp3-IRES-GFP (Foxp3-GFP) knock-in mice were kindly given by Dr Bernard Malissen (CIML, Marseille). Mice were housed under specific pathogen-free conditions. All experimental protocols were approved by the local ethics committee and are in compliance with European Union guidelines.

### Mass Cytometry

For labelling of antibodies, Maxpar Antibody Labeling Kit (Fluidigm) was used according to the manufacturer instructions. Briefly, lanthanide was loaded with metal and antibody was reduced, in parallel, using TCEP solution (Thermo Scientific). Then, loaded lanthanide was added to reduced antibody before incubation for 90min at 37°C, Finally, the metal-conjugated antibody was diluted in 100uL in Antibody Stabilizer (CANDOR® Bioscience). Up to 5.10^6^ live cells were stained with Cell-ID Cisplatin (Fluidigm) for 10min at RT. Fc receptor were blocked with anti-CD16/32 (clone 2.4G2) for 10min before staining with extracellular antibodies in PBS 1X Ca-Mg-for 25min at 4°C. Before intracellular staining, cells were fixed using 2% para-formaldehyde for 15min at 4°C. Then, True Nuclear Transcription Factor Buffer set (Biolegend) was used and cells were incubated with antibodies for intracellular staining during 40min at 4°C. Finally, cells were incubated with Cell-ID Intercalator-Ir (Fluidigm) in 2% PFA for 16 hours before freezing at −80°C until acquisition on Helios cytometer and CyTOF software version 6.0.626 (Fluidigm) at the Cytometry Pitié-Salpétrière core (CyPS). Dual count calibration, noise reduction, cell length threshold between 10 and 150 pushes, and a lower convolution threshold equal to 10 were applied during acquisition. Data files were normalized with the MatLab Compiler software normalizer using the signal from the 4-Element EQ beads (Fluidigm) as recommended by the software developers. To normalize the variability between mice for supervised (i.e 2D plots) and unsupervised (i.e tSNE) analysis, cell samples from 3 mice were pooled before stainings.

### Generation of tSNE plots

tSNE plots were generated with FlowJo v10.6.1 using 1000 iterations, a perplexity of 30 and an eta of 665 for the periphery and 881 for the tumor. The clustering was done in manually gated CD45+ cells with CD3, CD4, CD8a, Foxp3, B220, IA/IE, TCRb, CD11b, CD11c, Ly6G, Ly6C and NK1.1 as tSNE parameters. Immune subsets were defined as follow: B cells, B220+IA/IE+CD11b-CD3-; CD4 T cells, CD3+CD4+TCRb+; CD8 T cells, CD3+CD8+TCRb+; gamma-delta T cells, CD3+NK1.1-TCRb-; Monocytes/Macrophages, CD11b+Ly6G-NK1.1-; Neutrophils, CD11b+Ly6G+; NK cells, NK1.1+CD3-; NKT cells, NK1.1+CD3+

### Induction of EAE

To induce EAE, C57BL/6J mice were injected subcutaneously in the flanks with 100 μg of MOG35-55 peptide (Polypeptide) emulsified in 100 μl of CFA (Sigma-Aldrich) supplemented with 50 μg of heat-killed Mycobacterium tuberculosis H37Ra (BD Biosciences). Animals were additionally injected intravenously with 200 ng of Bordetella pertussis toxin (Enzo) at day 0 and 2 of EAE induction.

### Tumor models

For CyTOF analysis, mycoplasma-free MC38 (colon carcinoma) cells were grown in RPMI media supplemented with 10% FCS, L-glutamine and antibiotics (Penicillin/Streptomycin). 13 weeks-old C57/BL6J mice were injected subcutaneously with 5.10^5^ tumor cells or PBS 1X. At day 26, mice were sacrificed and spleen, axillary, brachial and inguinal lymph nodes and tumors were harvested. For RNA-Seq experiments, a similar protocol was used except that the MCA fibrosarcoma cell line was used.

### Tissue preparation and cell sorting

For CNS analysis, the spinal cord and the brain were harvested from mice. CNS samples were digested in type IV collagenase (1 mg/ml) and DNase I (100 μg/ml) for 30 min at 37°C. Tumors and peripheral lymphoid organs were digested with 0.84mg/mL of collagenase IV and 10μg/mL DNAse I (Sigma Aldrich) for 40min at 37°C with an intermediate flushing of the tissue. To eliminate dead cells and debris, cell suspensions were isolated on a Percoll gradient (40/80). Rings were collected, washed, and cell pellets were resuspended in PBS 3% SVF before counting. Tregs were purified after enrichment of CD25+ cells using biotinylated anti-CD25 mAb (7D4) and anti-biotin microbeads (Miltenyi Biotec), followed by CD4 staining (RM4.5) and cell sorting using the BD FACSAria II. Tconvs were purified after enrichment of CD25-cells using biotinylated anti-CD25 mAb (7D4) or of CD8-CD19-CD11b-cells using biotinylated anti-CD8 (53-6.7), CD19 (1D3) and CD11b (M1/70) mAbs and anti-biotin microbeads (Miltenyi Biotec), followed by CD4 staining (RM4.5) and cell sorting using the BD FACSAria II.

### TNFR agonists in vitro stimulation

The protocol has been described in details elsewhere (30). Briefly, purified Treg were sorted from Foxp3-GFP mice and 1,5.105 cells were cultured in 96-flat-well plates with anti-CD3/anti-CD28 (both coated at 2mg/ml) and IL-2 (10ng/ml). The following soluble TNFRSF agonists were used to co-stimulate Tregs: anti-4-1BB mAb (10 μg/ml, 3H3, BioXcell), OX40L (100 ng/ ml, AdipoGen), and the TNFR2 agonist TNC-sc(mu)TNF80 (STAR2) (12 ng/ml).

### RNA-Seq analysis

RNA was extracted from sorted cells using the NucleoSpin RNA XS kit from Macherey-Nagel, quantified using a ND-1000 NanoDrop spectrophotometer (NanoDrop Technologies) and purity/integrity was assessed using disposable RNA chips (Agilent High Sensitivity RNA ScreenTape) and an Agilent 2200 Tapestation (Agilent Technologies, Waldbrunn, Germany). mRNA library preparation was performed following manufacturer’s recommendations (SMART-Seq v4 Ultra Low Input RNA Kit, TAKARA Bio). Final samples pooled library prep was sequenced on Nextseq 500 ILLUMINA with HighOutPut cartridge (2×400Millions of 75 bases reads), corresponding to 2 times 23 × 10^6^ reads per sample after demultiplexing. Poor quality sequences were trimmed or removed with Trimmomatic software (31) to retain only good quality paired reads. Star v2.5.3a were used to align reads on reference genome mm10 using standard options. Quantification of gene and isoform abundances were done with RSEM 1.2.28, prior to normalization on library size with DESEq2 bioconductor package. Absolute counts per million reads (CPM) were retrieved to quantify *Penk* transcript abundance in our experimental datasets.

## Results

### Analysis of the GEO Treg meta-signature

As a starting point, we searched for a common Treg signature (that is a list of genes differentially expressed in Treg compared to conventional T cells (Tconv)) across publicly available datasets retrieved from the GEO web site. We manually selected 8 datasets based on subjective criteria explicited below, and availability of the GEO2R analytical tool for the dataset. We deliberately limited the search to cells isolated from peripheral lymphoid tissues at steady state. Among the 8 datasets, 4 had been generated from lymph node cells, 3 from spleens and 1 from the bone marrow, 7 had been generated in C57BL/6 and one in BALB/c mice. Various ages and sexes were present in the datasets (range: 6-36 weeks-old). Various Treg sorting strategies were also used, from classical CD25^+^ sorting to isolation of GFP^+^ cells from Foxp3-GFP transgenic mice. So we believe that the 8 chosen datasets were representative of typical Treg profiles found in many studies. For each dataset, we generated a list of differentially expressed genes with a cutoff based on a false discovery rate inferior to 0.05 and a log2 fold change superior to 1. We then established a list of differentially expressed genes common to the 8 datasets: 58 genes were found to be common to all 8 datasets. To gain biological insights from this list of genes, we generated a network of putative protein-protein interactions (PPI), protein-DNA and genetic interactions, pathways, reactions, gene and protein expression data, protein domains and phenotypic screening profiles from the Treg meta-signature (TMS) using the Genemania Cytoscape application, linking nodes of the network based on a score aggregating experimental and literature-based interactions (27) (Figure 1A). We represent the mean fold-change as a color code and the numbers of neighbors connected to each node (degree) as the size of the node to highlight some noticeable facts: *Foxp3 and Il2ra* were the most differentially expressed genes and were also among the most connected genes to other members of the network, as expected from the sorting Treg strategy used to generate these datasets. The well described Treg marker *Ctla-4* was also highly differentially expressed and connected to other members of the network. Only *Tnfrsf4* (OX40) was more connected to the network but had a slightly lower fold change, much like *Tnfrsfr9* (4-1BB) and *Tnfrsf1b* (TNFR2). Genes with high fold-change but lower degree were *Itgae* (CD103), *Rgs1*, *Klrg1*, *Ikzf2* (HELIOS), *Nrp1* and surprisingly *Penk*, a gene coding for the proenkephalin, precursor of Met-enkephalin, an important mediator of peripheral nociception (pain perception). This *in silico* isolation of many known Treg markers confirms the validity of our approach and suggest that novel molecules isolated by this method, such as *Penk*, could be considered as reliable Treg “markers”.

**Figure 1:**
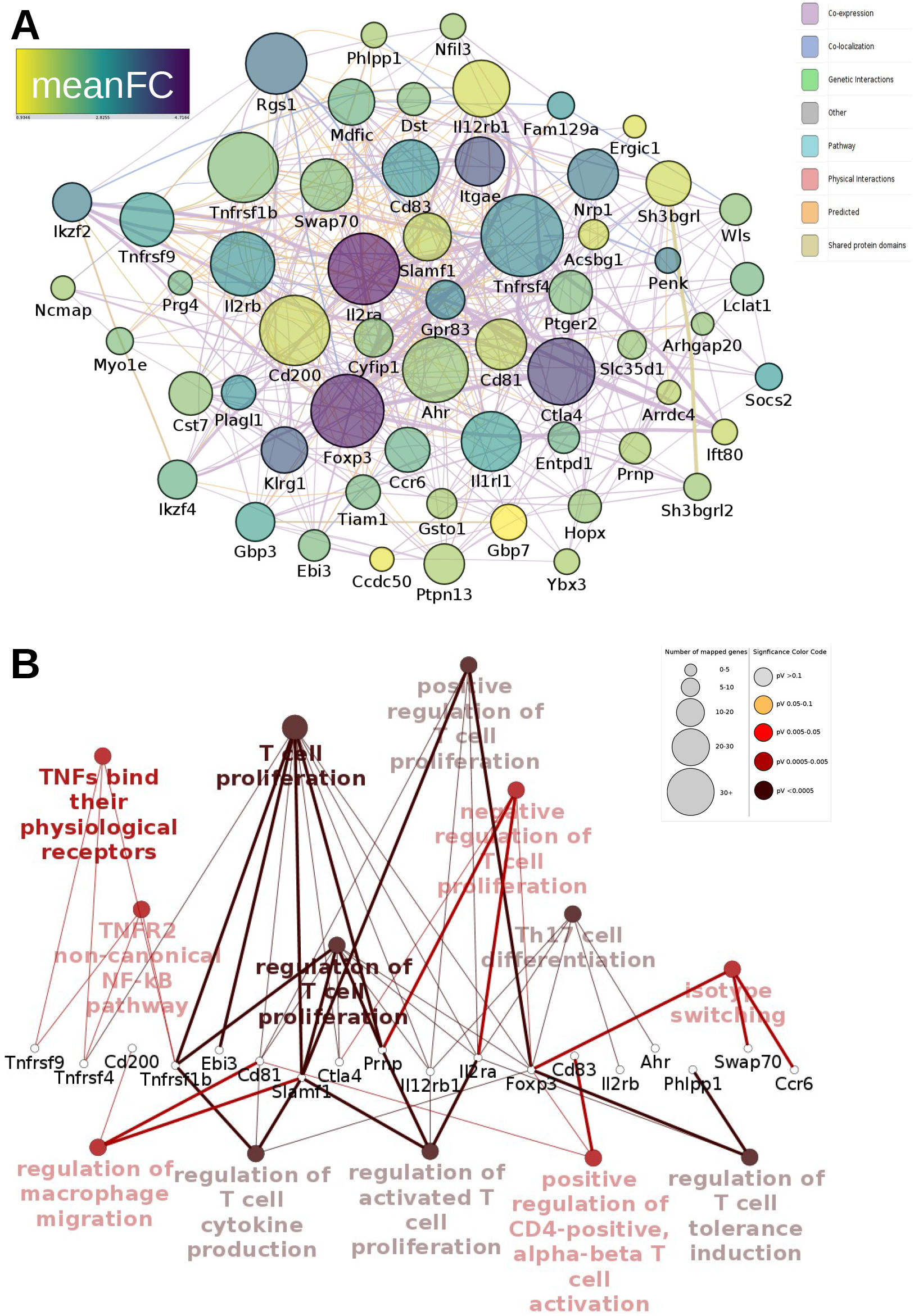
Analysis of the GEO Treg meta-signature. **(A)** The size of the node is proportional to the degree (number of connections to other nodes) of each node. The color indicate the mean fold change between Tconv and Treg across the 8 datasets (range: 0.9-4.7). The edge color (link between nodes) indicate the nature of the interaction, as detailed in the legend. Interaction network of the Treg-meta-signature (TMS) was generated with Genemania in Cytoscape. **(B)** Listed are the genes of the TMS common to different pathways and their associated ontology. The size of each node is proportionnal to the number of genes enriched in that node and the color indicate the statistical significance of the enrichment (Benjamini-Hochberg correction), as detailed in the legend. Enrichment analysis of the TMS was performed with ClueGO/Cluepedia in Cytoscape.

To gain further biological insights from the TMS, we performed an enrichment analysis using the ClueGO plugin in Cytoscape which interrogates several ontology/pathways databases simultaneously and produce a network of ontologies/pathways terms according to the statistical significance of the enrichment (26) (Figure 1B). As expected, most of the significant GO/pathway terms revolved around T cell proliferation and its regulation. Among the TMS, only few genes belong to several pathways at the same time and *Foxp3* was the most prominent one. Surprisingly, *Slamf1* (CD150) and *Prnp* (CD230) were also connected to several related ontologies. Of note is that TNF-related ontologies were enriched given the presence of 3 members of the TNF Receptor Superfamily in the TMS.

### Analysis of the Immuno-Navigator Treg meta-signature

To confirm and extend the findings, we then turned to the Immuno-Navigator database, a batch-corrected collection of RNA quantification across numerous studies and many samples and cell types in mice and humans (32). We compare 634 samples of CD4 T cells, that include many types of CD4+ T cell subsets with 240 samples of purified Treg, also defined in various ways and from various origins, including gene-deficient animals. Once variables with low variances were filtered out, a Principal Component Analysis (PCA) showed that the CD4 and Treg samples segregated well apart, with PC1 recapitulating 72% of the variance (Figure 2A), showing that the remaining variables in the datasets could discriminate the two subsets with high accuracy. This was also visible in the clustered heatmap, where Treg samples clearly segregated from CD4 samples (Figure 2B). A list of 56 up regulated genes in Treg filtered on the FDR and the fold change also establish a dense network of putative PPI, although many members were not connected (Figure supplemental 1). The topology of the network was very similar to the one of the TMS (Figure 1), with *Foxp3*, *Il2ra* and *Ctla4* being both highly differentially expressed and most connected to other members of the network. Of note is that *Cpe*, coding for Carboxypeptidase E, responsible with other convertases for processing proenkephalin into biologically active MENK peptides (33), was present in this signature. To go further, we next investigated the overlap between the Immuno-Navigator signature and the TMS. There was a 50% overlap in genes differentially expressed in CD4 and Treg from the Immuno-Navigator database relative to our more stringent TMS (Figure 2C). A list of 31 common genes was then projected into a network of PPI to search for enrichment for some biological functions (Figure 2D). Most members of this list were expressed at the plasma membrane and were involved in signal transduction. Interestingly, the list contains well-known markers of Treg (*Foxp3*, *Il2ra*, *Ctla4*, etc) but also some unexpected genes, including *Penk, Cst7,* or *CD83*. Thus, we propose that this condensed network constitute a “core” set of genes that defines Treg, and that *Penk* is a reliable Treg marker.

**Figure 2:**
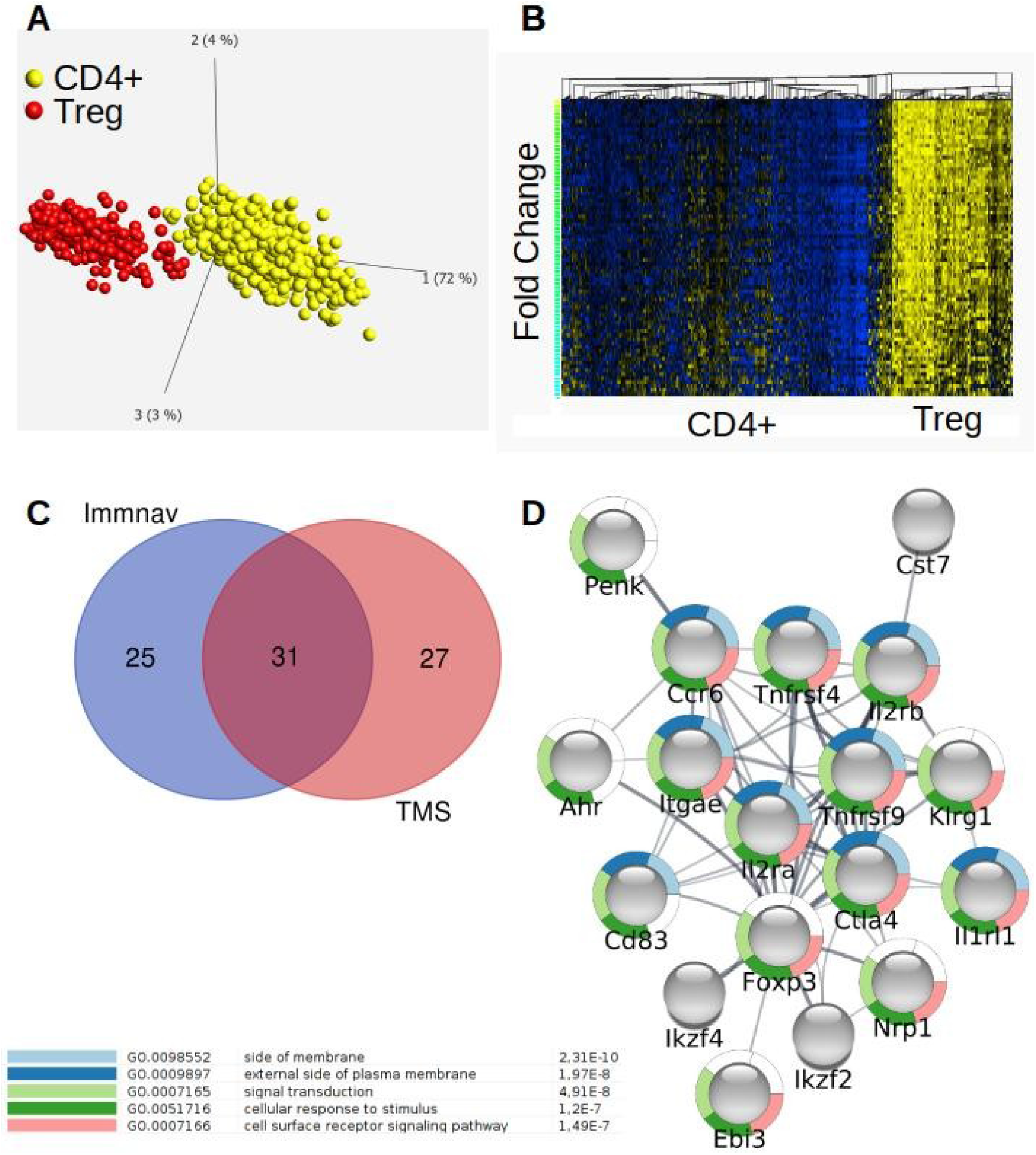
Analysis of the Immuno-Navigator Treg meta-signature. **(A)** Principal Component Analysis of CD4+ (yellow balls) and Treg (red balls) associated variables (mRNA transcript quantification) after filtration on the variance (0.5), the FDR (<0.05) and the fold change (>2). For more details on the structure of the Immuno-navigator dataset, please refer to (32). **(B)** Heatmap representing the z-score of differentially expressed genes in Treg relative to CD4+ cells with the same filters than in (A). Variables were sorted on the y-axis based on fold changes whereas samples were sorted by hierarchical clustering. Analysis in (A) and (B) were performed with Qlucore v3.4. **(C)** Venn diagram representing the intersection of the TMS with the Immuno-navigator Treg signature. **(D)** Putative Protein-Protein-Interaction network of the 31 common genes extracted from (C). The GO terms associated to each node is represented by a donut (color coded in the legend with the associated FalseDiscoveryRate). The PPI network was generated with STRING in Cytoscape.

### Mass cytometry analysis of the TMS

We next sought to determine whether this “core” set of genes was really restricted to Treg at the protein level relative to other immune cell subsets beyond Tconv. For that, we established the expression profile of some markers of the TMS in various immune cell subsets with a 38-parameters mass cytometry panel in peripheral lymphoid organs and in a syngeneic tumor model (Table supplemental 1). We manually defined the main immune subsets (CD4 and CD8 T cells, B cells, γδ T cells, NK and NKT cells and various subsets of myeloid cells) in gated CD45^+^ cells according to expression of lineages markers on a tSNE plot for the periphery (Figure 3A) and from the tumor (Figure 3B). In the periphery, a discrete cluster of FOXP3^+^CD25^+^CTLA-4^+^HELIOS^+^CCR6^+^4-1BB^+^CD103^+^ cells was visible in CD4^+^ T cells, presumably representing Treg. But in contrast to the first 4 molecules which were restricted to a subset of CD4+ T cells, CCR6 was also highly expressed by B cells, 4-1BB by a subset of NK cells and CD103 by a subset of CD8+ T cells, showing that these were not reliable Treg markers in the periphery. In tumor-infiltrating cells, a discrete cluster of FOXP3^+^CD25^+^CTLA-4^+^4-1BB^+^CD103^+^ could be observed. In contrast, HELIOS and CCR6 were expressed by other subsets, neutrophils and B cells/monocytes/macrophages, respectively. Of note is that Treg were positive for both KI67 and pRb, signing active proliferation in the periphery and the tumor (Figure Supplemental 2). Also, we observed that TNFR2, a member of the TMS, poorly segregated in Treg at the cellular level beyond the comparison with Tconv. Finally, other proteins known to mark Treg but not present in the TMS, such as PD-1 or ICOS, were also over expressed by Treg in the periphery. However, PD-1 was not a reliable Treg marker in the tumor whereas ICOS was highly expressed by Treg in the tumor. Likewise, GITR efficiently marked Treg in the periphery and in the tumor (Figure Supplemental 2) despite being absent from the TMS. Thus, among the proteins of the TMS that were investigated, only FOXP3, CD25, and CTLA-4 truly defined Treg across various immune subsets in the periphery and the tumor. Moreover, some proteins that were not in the TMS, such as GITR or ICOS, were restricted to Treg in the periphery and the tumor, highlighting the limitations of translating molecular data to the protein level.

**Figure 3:**
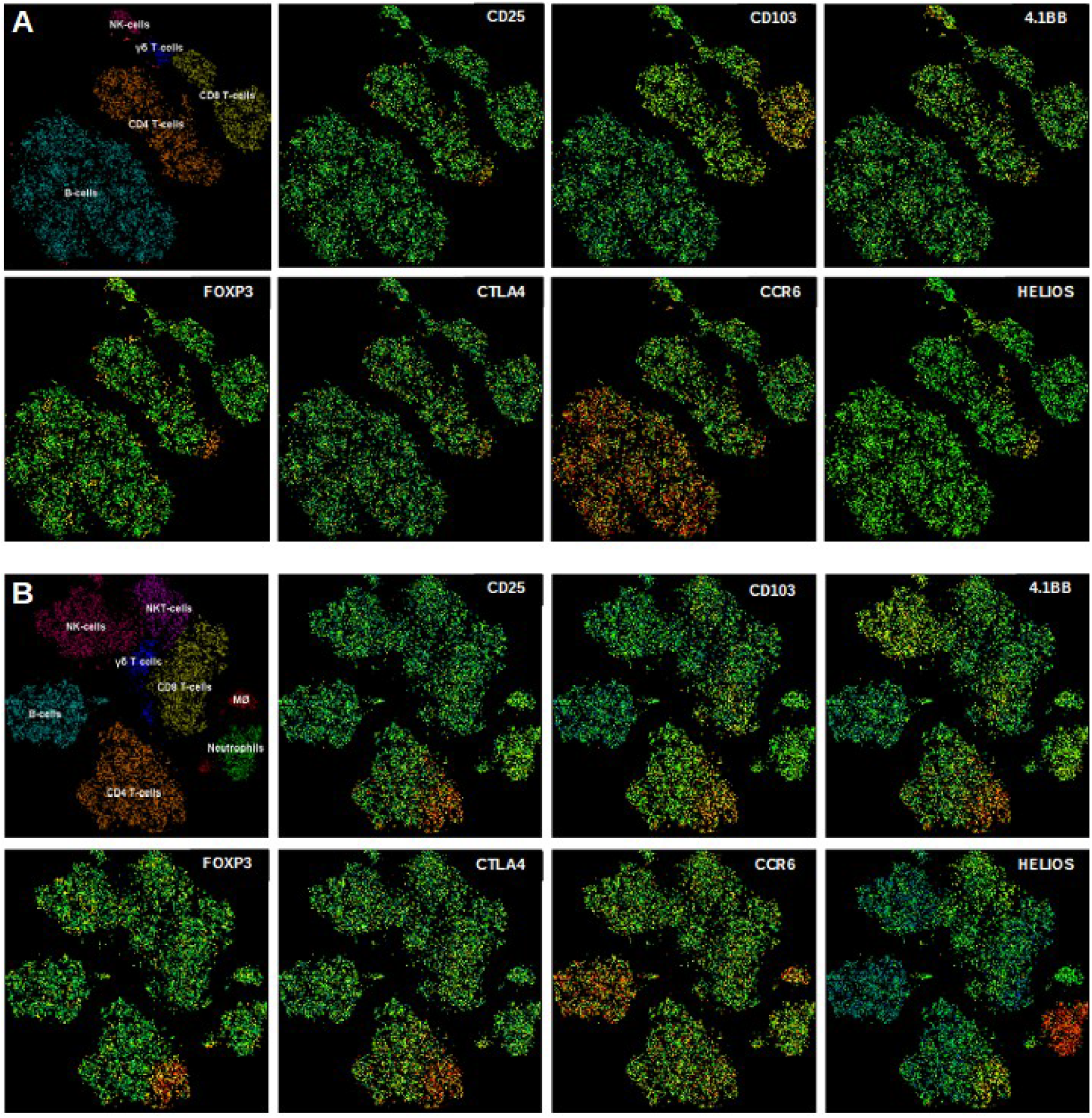
Mass cytometry analysis of the TMS. **(A)** Immune cell subsets identification on a tSNE plot gated on murine CD45^+^ cells in peripheral lymphoid organs and the projected expression of the indicated molecules. **(B)** Same as in (A) for the tumor tissue. Expression levels of each protein are depicted from green (negative to low) to red (moderate to high) except in the top left panels in A and B.

### Penk expression by Treg in various tissues and inflammatory conditions

Following on the surprising observation that *Penk* was a highly specific Treg genetic marker, we wanted to confirm this observation at the protein level. Deceptively, PENK was not listed in 2 different databases reporting the proteomes of murine Treg and Tconv, preventing any differential expression analysis to be performed (34,35). Furthermore, a monoclonal antibody to murine PENK is currently not available, preventing direct detection by cytometry. Then, we asked whether *Penk* mRNA over expression could be limited to peripheral lymphoid organs, since most of the data analyzed so far were generated in spleens or lymph nodes. A rapid survey of studies examining gene expression in Tconv and Treg in various tissues showed that *Penk* was found to be over expressed by Treg in the thymus (21) and in several tissues, including fat (17,18), and muscle (22) (Figure 4A). We also independently verified over expression of *Penk* in Treg relative to Tconv in our own set of RNA-Seq data from lymph nodes of normal mice (Figure 4B). Finally, we investigated whether *Penk* mRNA could be modulated in Treg in various inflammatory conditions (Figure 4C). In the CNS of mice undergoing Experimental Autoimmune Encephalitis (EAE), we observed that *Penk* was down modulated compared to non-inflamed lymph nodes. In contrast, we observed massive up regulation of *Penk* mRNA in Treg of the tumor relative to the draining LN.

**Figure 4:**
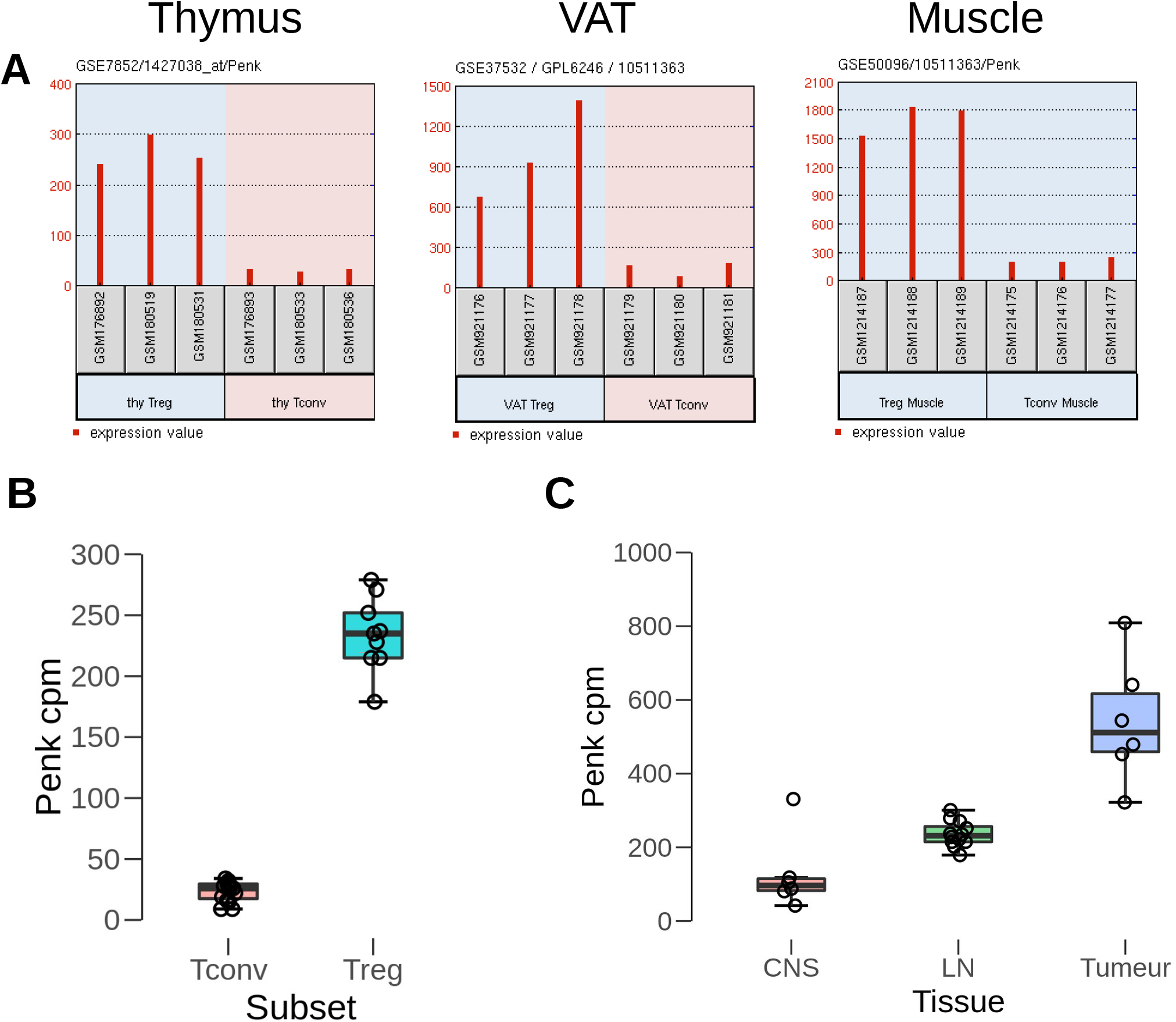
*Penk* mRNA expression in various tissues and in inflammatory conditions. **(A)***Penk* expression in Treg relative to Tconv in the thymus, Visceral Adipose Tissue (VAT), muscle in the indicated GSE datasets. Treg are highlighted in light blue, Tconv in light red. **(B)** RNA Seq analysis of *Penk* mRNA expression in Tconv and Treg from lymphoid organs of normal mice. **(C)** RNA-Seq analysis of *Penk* mRNA expression in Treg in the indicated conditions (CNS, central nervous system of EAE mice; LN, draining LN in EAE and tumor bearing mice, Tumor, Tumor-infiltrating Treg in the MCA model). Each dot is a mouse. EAE and tumor mice were different and their RNA-Seq was performed in different runs. No statistical analysis was performed given the large differences in the means between the groups.

### Analysis of the genes most correlated to Penk mRNA

In order to gain mechanistic insights into the regulation of *Penk* expression in Treg, we first established a network linking the genes most correlated to *Penk* mRNA from the Immuno-Navigator database in all cell types combined (Figure 5A). This network included several members of the TMS, including *Foxp3*, *Ikzf4*, *Ncmap*, *Ctla-4*, or *Gpr83*, showing that *Penk* was highly correlated with some genes of the TMS defined above. The high correlation between *Foxp3* and *Penk* is illustrated in figure 5B, where the co expression of *Foxp3* and *Penk* in Treg is also clearly visible. We next interrogated the Immuno-Navigator database to establish a network of the genes most correlated to *Penk* specifically in Treg (Figure 5C). *Penk* was highly correlated to 4 TNFR members (*Tnfrsf1b* (TNFR2), *Tnfrsf4* (OX40), *Tnfrsf9* (4-1BB) and *Tnfrsf18* (GITR)), and to the TF *Batf* (illustrated in figure 5D), suggesting that *Penk* expression in Treg might be regulated by TNFR signaling and BATF.

**Figure 5:**
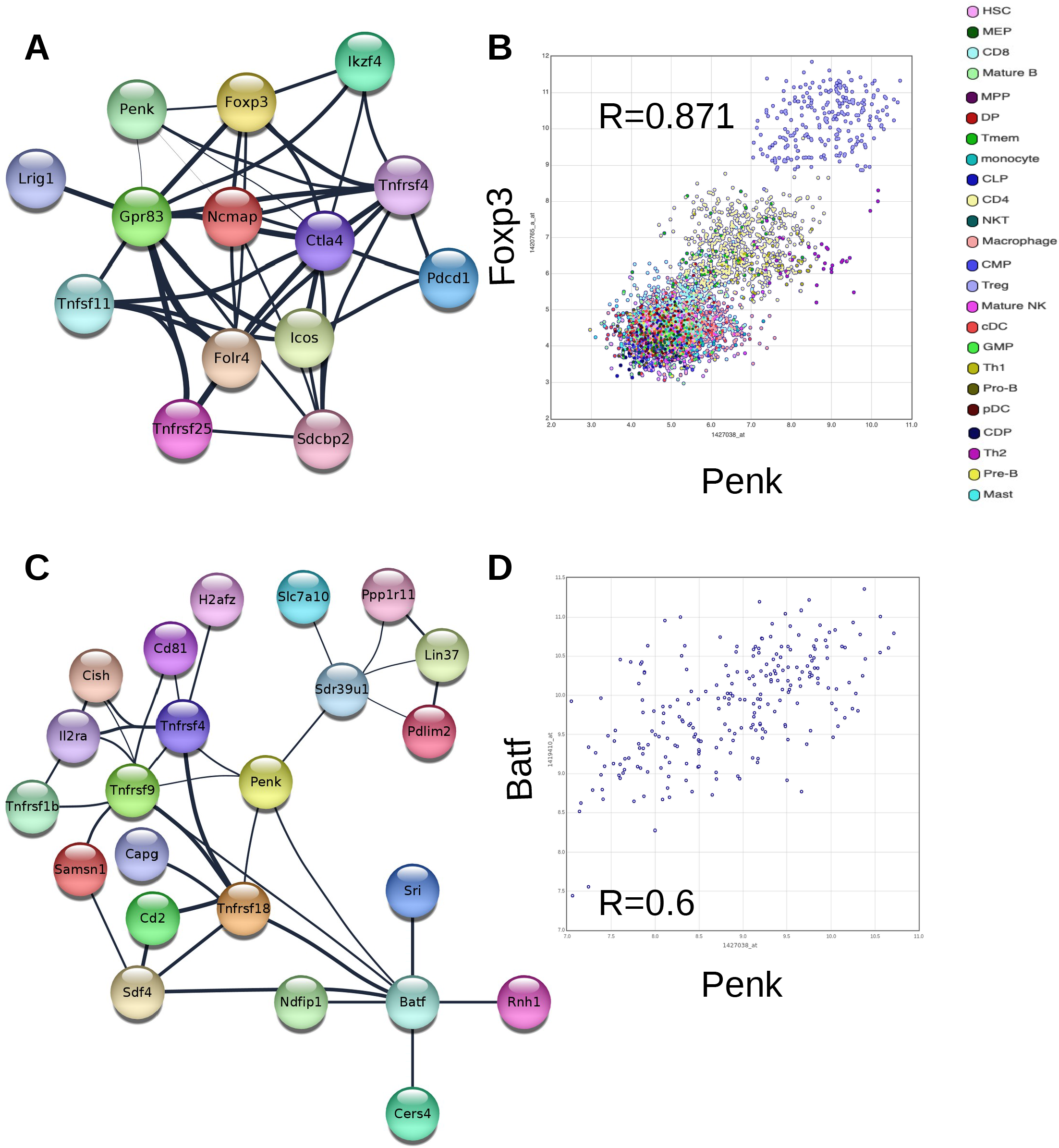
Analysis of the genes most correlated to *Penk* mRNA. **(A)** The genes most correlated to *Penk* in all cell types combined. The Pearson correlation values were extracted from the Immuno-Navigator database and integrated into Cytoscape. Each node is a gene linked by edges with width proportional to the Pearson correlation. All were greater than 0.7 (range: 0.774-0.936). **(B)** Illustration of the correlation between *Penk* and *Foxp3* in all cell types listed in the legend. **(C)** Same as in (A) for the genes most correlated to *Penk* in Treg only (edge range: 0.538-0.758). **(D)** Illustration of the correlation between *Penk* and *Batf* in Treg. Each dot is a sample from the Immuno-Navigator database.

### Penk mRNA expression is regulated by TNFR signaling and BATF-AP-1

To investigate this possibility, we turned to our own set of data establishing *in vitro* the transcriptome of TNFR-stimulated Treg by RNA sequencing (30). Indeed, *Penk* expression was increased relative to controls with TNFR2, OX40 and 4-1BB agonists and this was more marked at 36h post stimulation (Figure 6A). Furthermore, the expression of *Batf* was also increased by TNFR agonists relative to controls and at an earlier time point than *Penk* (Figure 6B), showing that up regulation of *Batf* preceded up regulation of *Penk*, consistent with a model in which the TF BATF positively regulated *Penk* mRNA transcription. A prediction of that model would imply a reduction in *Penk* mRNA in BATF-KO Treg. Indeed, GEO2R analysis of the transcriptome of BATF-KO Treg (36) showed a dramatic decrease in *Penk* expression relative to control Treg (Figure 6C). Finally, BATF was reported to control several immune regulatory networks through AP-1 members JUN or FOS (37). Interestingly, members of the AP-1 TF were listed as potential regulators of *Penk* mRNA according to the Ingenuity database (Figure 6D). Altogether, our results point to a model in which TNFR signaling regulates *Penk* expression in Treg through modulation of the BATF-AP-1 complex.

**Figure 6:**
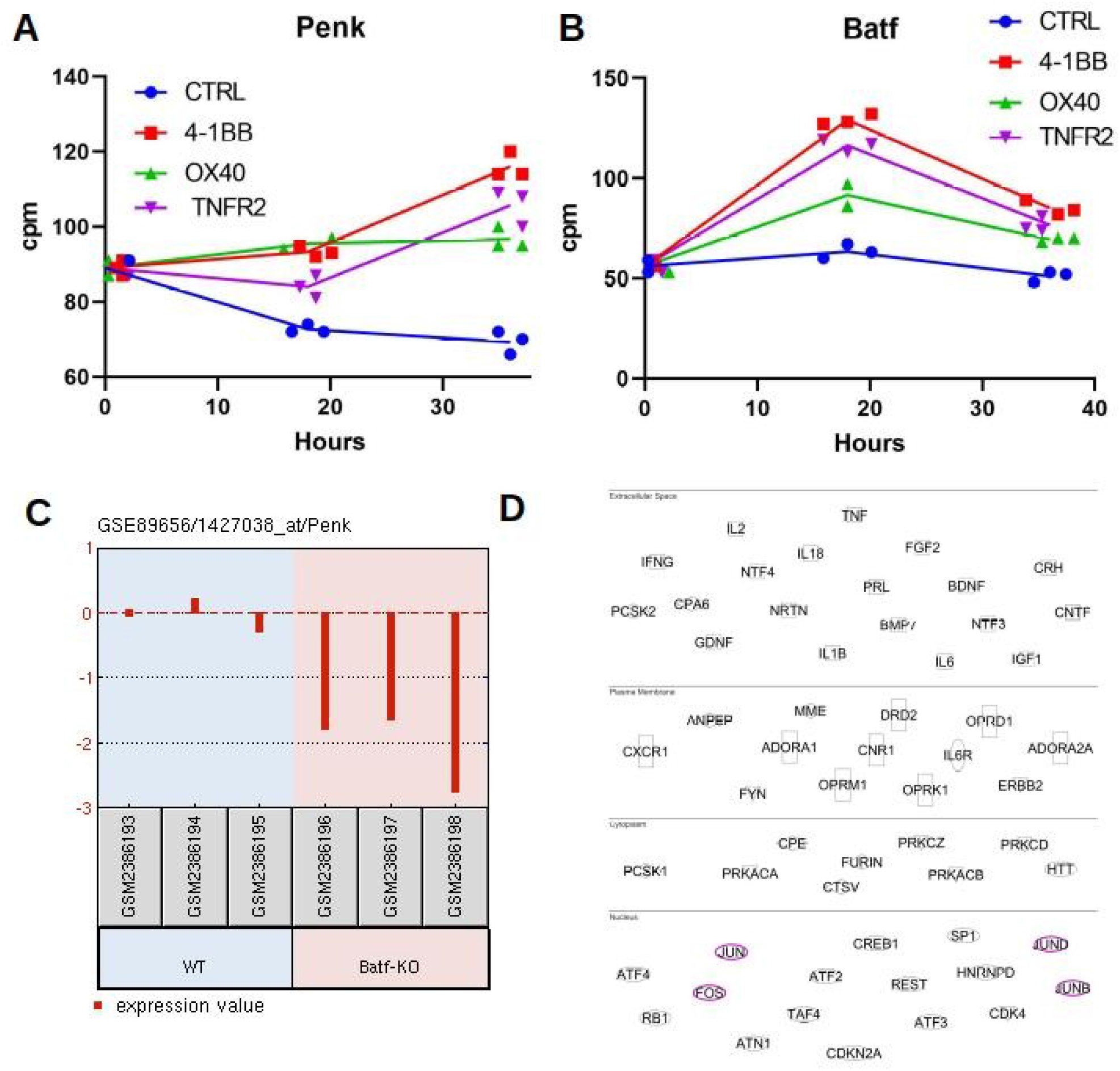
*Penk* mRNA expression is regulated by TNFR signaling and BATF-AP-1. *Penk* **(A)** and *Batf* **(B)** mRNA expression after in vitro stimulation of purified Treg with the indicated TNFR agonists prior (0), and at 18 and 36hrs after stimulation. Each dot is a biological replicate from a single experiment. **(C)** GEO2R analysis of the GSE89656 dataset for *Penk* mRNA variations between wild type control Treg (WT) and BATF-KO Treg. **(D)** Direct and indirect regulators of *Penk* mRNA were extracted from the Ingenuity Pathway Analysis database according to their cellular localization indicated in the figure. AP-1 TF are highlighted.

## Discussion

To our knowledge, the present report is the first attempt to define a “universal” Treg signature in mice by a meta analysis of published and available datasets. Our TMS provide a more solid appreciation of Treg-specific genes relative to the Immuno-navigator database which includes many types of heterogeneous samples. Our unbiased approach confirms that the genes belonging to the interleukin-2 signaling pathway (*Il2ra*, *Il2rb*) and of the TNF receptor super-family (*Tnfrsf4, Tnfrsf9*) are central to their identity. Our gene ontology enrichment analysis of the TMS also confirms that the “core” set of genes belongs to biological processes important for Treg function; regulation of T cell proliferation and cytokine signaling pathway. Finally, the preferred cellular localization of the TMS gene products at the cell surface highlights the dependency of Treg to external stimulus for their function and to preserve their identity. This is a view supported by functional plasticity of Treg (effector class control), and possible acquisition of effector functions in some inflammatory conditions. One point of caution for the interpretation of PPI networks is that mRNA expression may or may not be correlated with protein expression, and thus with biological functions. For instance, a recent study has found strong discrepancies between mRNA and protein levels in human Treg (38). This low correlation between mRNA and protein levels was also observed in human Treg induced in vitro (39) but surprisingly not in murine “natural” Treg (34). Whether this is a true specie-specific difference or whether it is related to technical issues remains to be investigated. This is an important matter since it relates to the more general question of inferring biological insights from a list of differentially expressed genes. Collecting a proteomic Treg meta-signature from all available proteomic studies in mice, properly normalized and corrected for batch effect, will bring essential informations to that matter.

A few genes stand out to be highly enriched in Treg relative to other immune subsets at the protein level and some significant differences were noted when the periphery and the tumor were compared. Everything combined, we established that, relative to Foxp3^+^ cells, CD25, CTLA-4, 4-1BB, GITR and ICOS were the most reliable Treg markers in our panel. It is interesting to note that some of those were not present in the TMS, indicating that they may have been filtered out during differential expression analysis or that their mRNA expression does not faithfully reflect their expression at the protein level, as described above. Nevertheless, these molecules and their ligands should probably concentrate the efforts for therapeutic targeting of Treg. The former is already in the clinics (Ipilimumab) with great efficacy to affect Treg but with possible severe adverse events whereas 4-1BB agonists are currently being tested in clinical trials (40). IL-2 has been a long standing history for cancer immunotherapy but its propensity to enhance Treg function makes it a difficult candidate to use as is. Based on our observations, we suggest that combination therapy targeting CD25, GITR, ICOS and/or 4-1BB might represent the most effective mean to affect Treg preferentially over Tconv while leaving other immune cells untouched. Indeed, intra-tumoral Treg depletion by monoclonal antibodies to CD25 (9), GITR (41), 4-1BB (42) or ICOS (43) was shown to be an efficient strategy for cancer immunotherapy.

A surprising and serendipitous finding in the quest for a “universal” Treg signature was the presence of the *Penk* gene in the “core” set of genes defining Treg. Enriched *Penk* expression by Treg has been reported before in Treg clones derived from TCR-transgenic mice (44) and in brain Treg of mice recovering from stroke (45). Our analysis significantly extends these observations by showing that *Penk* expression by Treg is consubstantial of their generation in the thymus, independent of their localization, and can be modulated in the tissues during inflammation. Based on our own and previously published datasets, we propose a model in which TNFR signaling regulate *Penk* mRNA transcription through modulation of the Batf/AP-1 complex. Interestingly, *Penk* was down modulated in Treg in the EAE model, where Treg are defective, whereas it was up regulated in Treg of the tumor, where Treg are suppressors, suggesting an immunosuppressive role for Penk-derived opioid peptides. This has been clearly established *in vitro* on PHA-activated murine T cells (46). Accordingly, EAE was less severe after *in vivo* injection of the MENK peptide and that correlated with reduced numbers of T cells in the draining lymph nodes (47,48). However, mice deficient for PENK were also protected from EAE, in contradiction with a protective role for MENK in EAE (49). These discrepancies might be explained by the fact that PENK is also expressed by cells of the CNS with unknown impact on the development of the disease. To our knowledge, there is to date no studies looking for a role of immune-derived MENK on EAE physiopathology. Similar hypothesis for an immunosuppressive role of MENK was drawn from studies in another autoimmune setting, namely colitis induced by transfer of PENK-sufficient or deficient T cells in immunodeficient hosts (50). Finally, a modest impact of MENK peptides on tumor growth has been reported to be mediated by dendritic cells and possibly Treg (51). To date, there is to our knowledge no studies assessing tumor growth in animals deficient for PENK only in the immune system, precluding any firm conclusions on the role of endogenous MENK to be drawn. Based on enriched *Penk* expression in Treg, we would like to speculate that peripheral nociception might be intermingled with immune regulation at the Treg level, a novel hypothesis testable in wet lab experiments. It will be particularly crucial to analyze mice specifically deficient for *Penk* in Treg to determine the positive or negative impact of *Penk* on the immune response and how that relates to peripheral nociception.

## Abbreviations

Treg: Regulatory T cells
Tconv: Conventional T cells
TNFRSF: Tumor Necrosis Factor Receptor Super Family
TMS: Treg Meta-Signature
TF: Transcription Factors
Penk: pro-enkephalin
MENK: Met-Enkephalin
PPI: Protein-Protein Interactions
FDR: False Discovery Rate
EAE: Experimental Autoimmune Encephalitis

## Acknowledgments

The author wish to thank Dr J Divoux for performing the MCA tumor experiments for RNA-Seq analysis, Dr Maryam Khosravi for performing the EAE experiments for RNA-Seq experiments, Dr Martina Lubrano di Ricco for performing the TNFR agonists for RNA-Seq experiments, Dr Armanda Casrouge and Mr Claude Baillou for cell sorting, Dr Béhazine Combadière for access to Ingenuity Pathway Analysis software, Dr Aurélien Corneau for running the CyTOF, all members of the ITAC team and all the members of the CIMI-PARIS research center for their encouragements and criticisms, and the curators of the different databases used in this study.

