## Supplementary figures and images for "Characterization of a regulatory T cells molecular meta-signature identifies the pro-enkephalin gene as a novel marker in mice"

### Figure supplemental 1

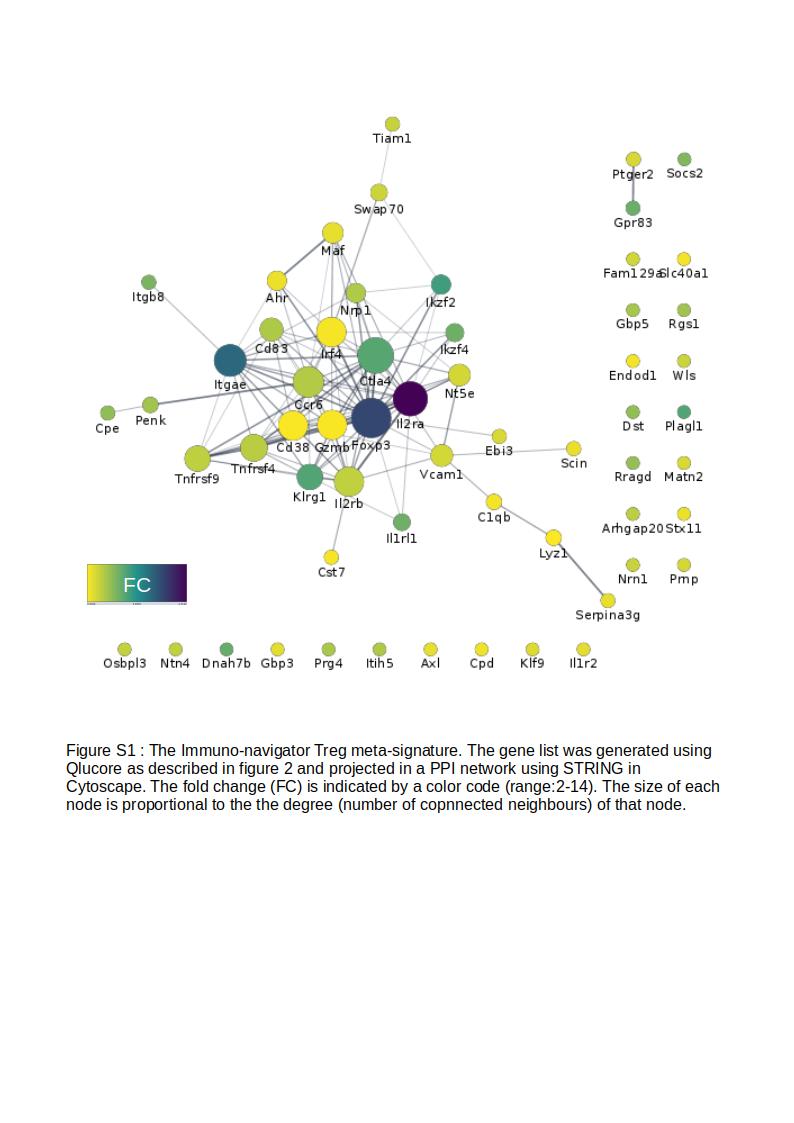

### Figure supplemental 2

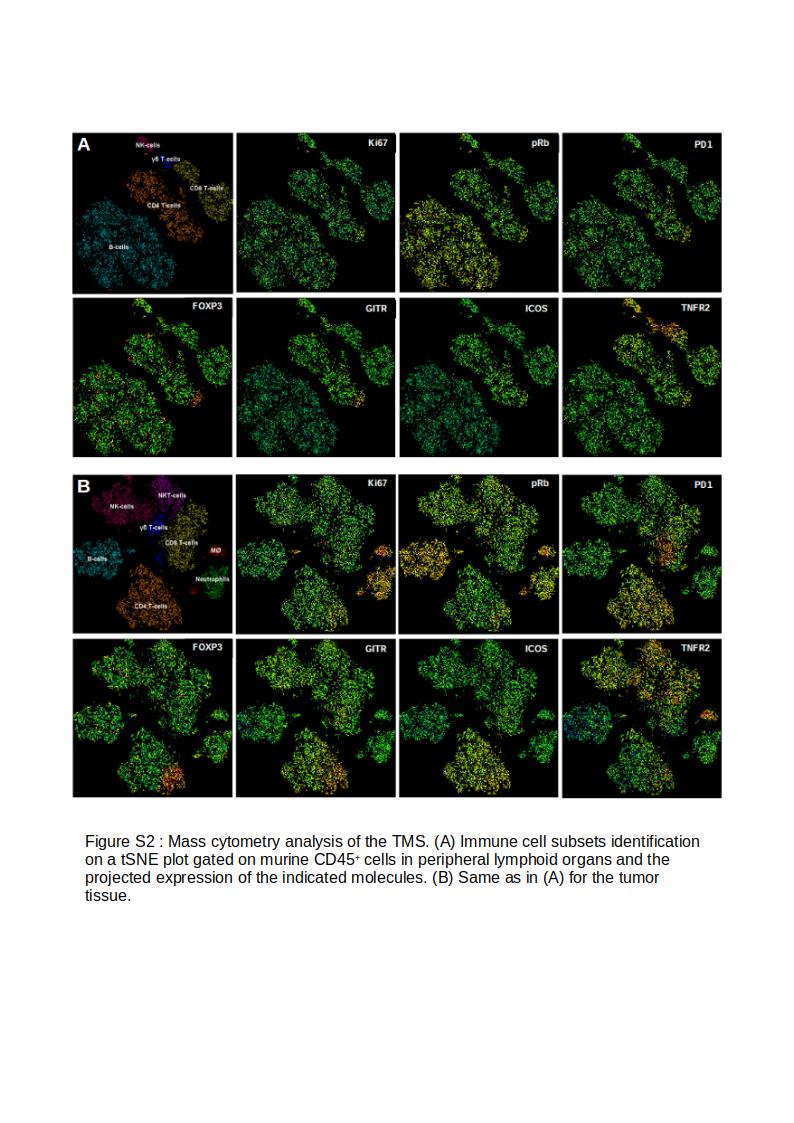
