## Supplementary material for "Characterization of a regulatory T cells molecular meta-signature identifies the pro-enkephalin gene as a novel marker in mice": Table supplemental 1

| Metal | Target | Clone | Distributor |
| --- | --- | --- | --- |
| 89Y | CD45 | 30-F11 | Fluidigm |
| 141Pr | Ly6G | 1A8 | Fluidigm |
| 142Nd | CD11c | N418 | Fluidigm |
| 143Nd | GITR | DTA-1 | Fluidigm |
| 144Nd | B220 (CD45R) | RA3-6B2 | Fluidigm |
| 145Nd | CD69 | H1.2F3 | Fluidigm |
| 146Nd | CD8a | 53-6.7 | Fluidigm |
| 147Sm | CD36 | NO.72-1 | Fluidigm |
| 148Nd | CD11b | M1/70 | Fluidigm |
| 149Sm | 4.1BB | LOB12.3 | BioXcell |
| 151Eu | CD25 | 3C7 | Fluidigm |
| 152Sm | CD3 | 145-2C11 | Fluidigm |
| 153Eu | CD206 | C068C2 | Biolegend |
| 156Gd | CCR6 | 29-2L17 | Fluidigm |
| 159Tb | PD1 | 29F.1A12 | Fluidigm |
| 162Dy | Ly6C | HK1.4 | Fluidigm |
| 163Dy | TNFR2 | TR75-54.7 | BioXcell |
| 164Dy | CD62L | MEL-14 | Fluidigm |
| 165Ho | NK1.1 | PK136 | Fluidigm |
| 169Tm | TCRb | H57-597 | Fluidigm |
| 170Er | CD183 | CXCR3-173 | Biolegend |
| 171Yb | CD44 | IM7 | Fluidigm |
| 172Yb | CD4 | RM4-5 | Fluidigm |
| 174Yb | CD103 | M290 | BD pharmingen |
| 175Lu | OX40 | OX86 | BioXcell |
| 176Yb | ICOS | 7E.17G9 | Fluidigm |
| 209Bi | IA/IE | M5/114.15.2 | Fluidigm |
| 150Nd | pRb | SER807/811 | Fluidigm |
| 154Sm | CTLA4 | UC10-4B9 | Fluidigm |
| 155Gd | RORgt | Q31-378 | BD pharmingen |
| 158Gd | Foxp3 | FJK-16S | Fluidigm |
| 160Gd | TBet | 4B10 | Fluidigm |
| 161Dy | iNOS | CXNFT | Fluidigm |
| 166Er | Helios | 22F6 | Biolegend |
| 167Er | Gata3 | TWAJ | Fluidigm |
| 168Er | Ki67 | B56 | Fluidigm |
| 173Yb | GzB | GB11 | Fluidigm |

Supplemental Table 1 : mass cytometry panel used in figure 3. Indicated are the metals coupled with the antibodies, the clones and the distributors. Antibodies not from Fluidigm were labelled in house.
